# Effects of benzothiazinone and ethambutol on the integrity of the corynebacterial cell envelope

**DOI:** 10.1101/2023.10.04.560928

**Authors:** Fabian M. Meyer, Urska Repnik, Ekaterina Karnaukhova, Karin Schubert, Marc Bramkamp

**Affiliations:** Institute for General Microbiology, Christian-Albrechts-University Kiel, Am Botanischen Garten 1-9, 24118 Kiel, Germany; Central Microscopy Facility, Christian-Albrechts-University Kiel, Am Botanischen Garten 1-9, 24118 Kiel, Germany; Faculty of Biology, Ludwig-Maximilians-University Munich, Großhaderner Straße 2-4, 82152 Planegg-Martinsried, Germany

**Keywords:** *Corynebacterium glutamicum*, arabinogalactan, mycolic acids, elongasome, benzothiazione, ethambutol

## Abstract

The mycomembrane (MM) is a hydrophobic layer formed by mycolic acids covering the surface of *Mycobacteria* and related species. This group includes important pathogens such as *Mycobacterium tuberculosis*, *Corynebacterium diphtheriae*, but also the biotechnologically important strain *Corynebacterium glutamicum*. The MM contributes to the impermeability of the cell envelope and thereby also protects bacteria from antibiotics. This makes biosynthesis of the MM an attractive target for antibiotic intervention. The first line anti-tuberculosis drug ethambutol (EMB) interferes with the synthesis of the arabinogalactan (AG), which is a structural scaffold for covalently attached mycolic acids that form the inner leaflet of the MM. Similarly, the new drug candidate, benzothiazinone 043 (BTZ) affects the synthesis of the AG component of the cell wall. We previously showed that *C. glutamicum* cells treated with a sublethal concentration of EMB lose the integrity of the MM. In this study we examined the effects of a sublethal concentration of BTZ. Our work shows that 1 µg ml^−1^ BTZ efficiently blocks the apical growth machinery and reduces cell proliferation, however the integrity of the MM is largely preserved and the effects of β-lactam antibiotics are only additive, not synergistic. Transmission electron microscopy (TEM) analysis revealed a distinct middle layer in the septum of control cells considered to be the inner leaflet of the MM covalently attached to the AG. It functions as a greasy slide for the lateral flow of mycolic acids in the outer leaflet. This layer was not detectable in the septa of BTZ or EMB treated cells, which suggests that the greasy slide is impaired and the confluency of the MM is thereby reduced. In addition, we observed that EMB treated cells have a thicker and less electron dense peptidoglycan (PG) layer consistent with the report that EMB also inhibits the glutamate racemase MurI. We conclude that EMB and BTZ have distinct differences in their modes of action. While EMB and BTZ both effectively block elongation growth, BTZ also strongly reduces septal cell wall synthesis. This renders BTZ treated cells likely more tolerant to antibiotics that act on growing bacteria.

## Introduction

Tuberculosis (TB) is the bacterial infection causing most annual fatalities (WHO, 2021). Consequently, its causative agent, *Mycobacterium tuberculosis* (MTB) is subject to intensive research with a particular aim to understand molecular mechanisms of infection. An increasing problem to treat TB infections is the spreading of antibiotic resistance within MTB strains. The emergence and spread of specific, and often concomitant, resistances against widely used drugs resulted in the propagation of multiple drug resistant (MDR) and extensively drug resistant (XDR)-MTB strains (Klaos et al., 2022; Seung et al., 2015; Sreevatsan et al., 1997; Telenti et al., 1993; WHO, 2021; Zhang et al., 1992). The spread of MDR and XDR strains impaired medical treatment regimens that have been established with existing anti-TB drugs. This prompted an intensive search for new anti-TB drugs in recent years. As a consequence, several new compounds that are potentially suitable for TB chemotherapy were described. One of the most recent additions to this list, after its US approval in 2012, was the mycobacterium specific ATPase-inhibitor bedaqueline (Hards et al., 2015; Kakkar and Dahiya, 2014). Currently two dozen of candidate substances are being clinically tested, but only four are subject to market approval (Stop TB Partnership, 2023). Among the list of anti-TB substances that are currently evaluated in clinical trials phase II, benzothiazinone 043 (BTZ) represents a new class of narrow bandwidth antibiotics that is reported to act on the synthesis of the arabino-galactan (AG) cell wall polymer (Batt et al., 2012; Makarov et al., 2009). This AG layer represents the connecting structure between the peptidoglycan (PG) sacculus and the outfacing, hydrophobic mycomembrane (MM) (Fig. 1).

**Figure 1:**
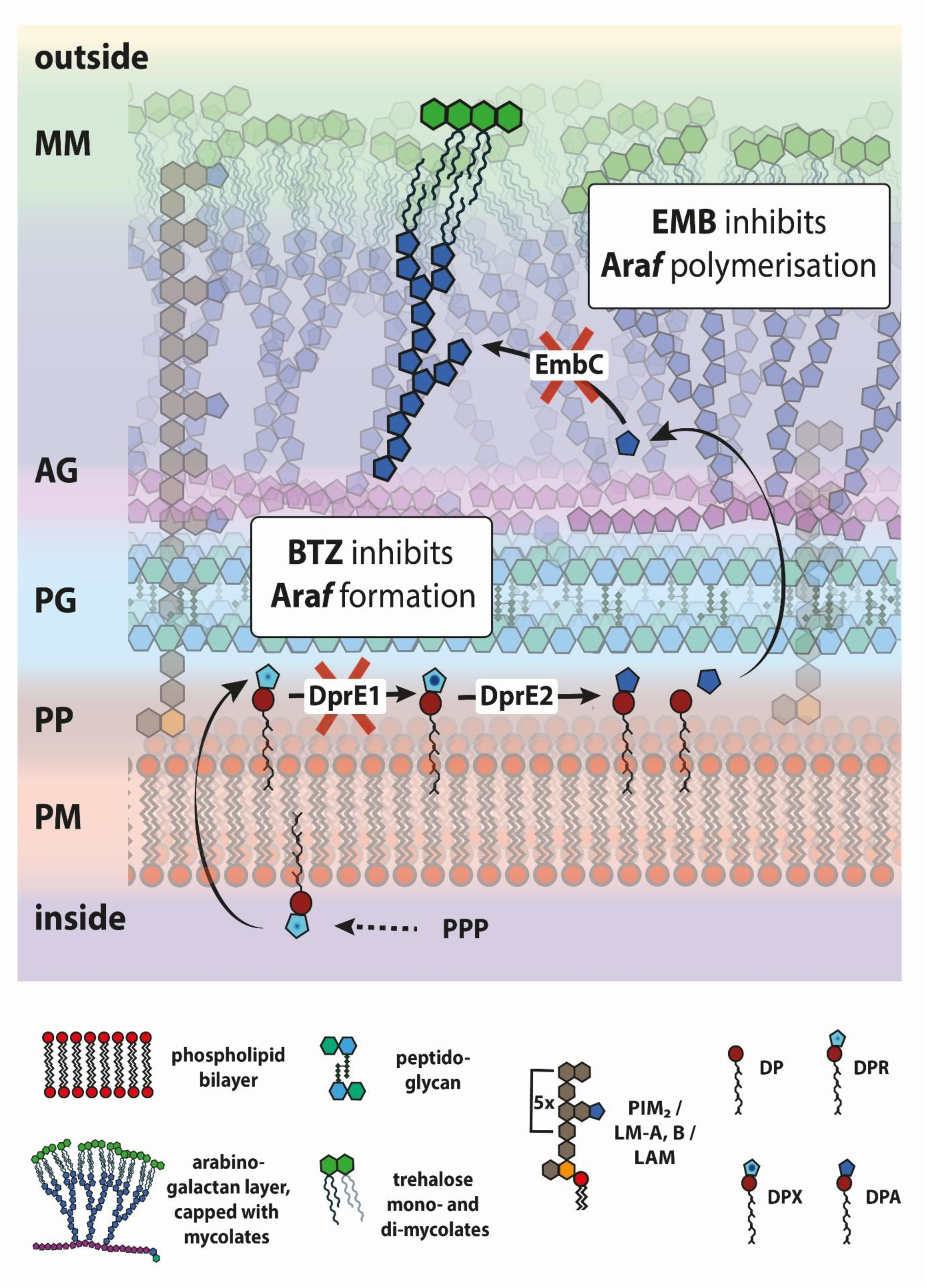
Cell envelope of *C. glutamicum*. Model of the corynebacterial cell that includes the major structural features of the cell envelope: mycomembrane (MM, green), arabinogalactan (AG, blue/purple), peptidoglycan (PG, lightgreen/cerulean), periplasm (PP), plasma-membrane (PM, red), scaffolding lipoarabinomannan species (grey). Further indicated are the sites of action for the front-line antituberculosis drug ethambutol (EMB) and benzothiazinones (BTZ), a novel drug substance currently under clinical trials. While EMB inhibits the polymerization of the AG by blocking paralogues of the enzyme Emb, BTZ interferes with the formation of the arabinose building blocks at an earlier stage by a suicide-inhibition of DprE1.

The molecular target of BTZ is the epimerase DprE1. The DprE1/2 complex that catalyzes a 2-step epimerization of decaprenyl-phospho-ribose (DPR) to decaprenyl-phospho-arabinose (DPA), a key precursor that serves as the prenol-linked arabinose donor required for the synthesis of cell-wall arabinans (Trefzer et al., 2012). Lack of arabinose synthesis by DprE1 inhibition impairs the formation of the arabinofuranose (Ara*f*) building blocks in the periplasm (Fig. 1) (Brecik et al., 2015; Grover et al., 2014; Mikusová et al., 1995). The blocked epimerization thereby leads to the accumulation of DPR and to the depletion of the recycling pool of decaprenyl phosphate, which ultimately causes cell death (Grover et al., 2014) (Fig. 1).

In contrast to the good understanding of its biochemical mechanism, the effects of BTZ on cell biology of bacteria are less well understood. Homologues of DprE1 from *M. tuberculosis* are present in clinically relevant taxonomic relatives like *M. leprea*, *C. diphteriae*, *Nocardia farcinica,* and also in the non-pathogenic *C. glutamicum*. The common denominator of this CMN (Corynebacteria, Mycobacteria, and Nocardia)-group bacteria is a characteristic multilayered cell envelope that consists of the plasma membrane, the periplasm, the peptidoglycan (PG), the AG and the mycomembrane (MM) (Dulberger et al., 2020; Zuber et al., 2008) (Fig. 1) The MM is a hydrophobic surface layer that contributes to the impermeability of the cell envelope and thereby protects against (bio-)chemical threats and hostile environmental conditions (Barry, 2001; Belisle et al., 1997; Dover et al., 2004) It is composed mostly of mycolic acids (MA) that are arranged in two leaflets. In the outer leaflet, the freely diffusible MA are free or attached to trehalose sugar to make trehalose monomycolate (TMM) or trehalose dimycolate (TDM). In the inner leaflet, MA) are covalently attached to the underlying AG layer and form a stable greasy slide for the lateral flow of the outer layer (Hett and Rubin, 2008; Marchand et al., 2012; Mikusová et al., 1995; Zuber et al., 2008). The length of MA residues defines the degree of hydrophobicity and serve as a taxonomic trait within the CMN-group. In the case of *M. tuberculosis*, with MA length of up to 90 carbon atoms, colonies show a waxy, dry surface and individual cells stain acid-fast. In contrast, the MM of corynebacteria is built on 40-carbon MA, resulting in a moist colony-surface and negative acid-fast staining (Chiaradia et al., 2017; Collins et al., 1982; Yassin, 2011; Zhou et al., 2019). Advantages such as a fast generation time, work-safety, and a well-established collection of genetic tools make *C. glutamicum* an ideal model to study cellular effects of drugs that target the synthesis of this complex CMN envelope.

The antibiotic EMB, which targets the synthesis of the arabinogalactan layer by inhibiting the polymerization of arabinose residues, was shown to increase the permeability of a cell wall in *M. vaccase* to hydrophobic compounds thereby sensitizing cells to erythromycin and rifampicin antibiotics (Korycka-Machała et al., 2005; Rumijowska-Galewicz et al., 2008). We have previously shown that EMB impairs the confluency of the MM in *C. glutamicum* and suggested that this contributes to the synergistic effects of rifampicin and β-lactam antibiotics in combination with EMB (Schubert et al., 2017). Since BTZ, like EMB targets the synthesis of the AG, in the present study we characterized its effects on cell biology of *C. glutamicum* and compared them to the effects of EMB.

Using fluorescence microscopy, we found that like EMB, BTZ inhibits the apical peptidoglycan (PG) synthesis, thereby effectively blocking elongation growth. Compared to EMB, BTZ also blocks cell division more efficiently. However, the MM layer was impaired less by BTZ than by EMB as shown by fluorescence microscopic and scanning electron microscopic (SEM) analyses. Relative quantification of the MM via thin layer chromatographic (TLC) analyses from crude cell wall extracts and analysis of MM-dynamics via fluorescence recovery after photo-bleaching (FRAP) experiments showed only subtle differences between the control and BTZ treated cells. Transmission electron microscopy (TEM) analysis revealed further structural differences in the cell envelope between BTZ and EMB treated cells, which likely contribute to increased sensitivity for β-lactam antibiotics in combination with EMB, but not with BTZ treatment as shown by antibiotic synergy testing. Differences between the two antibiotics and control cells were most notable at septa. An electron translucent layer in the middle of the septum that likely corresponds to the inner leaflet of the MM was observed only in control cells. We propose that this layer forms a stable greasy slide that allows the outer leaflet of the MM to flow into the septa during daughter cell separation and to cover the young cell poles.

## Material & Methods

### Cultivation, growth, and cell-wall preparation

Strains used in this study are listed in Table 1. *Corynebacterium glutamicum* 13032 cells and the derivatives *C. glutamicum divIVA::divIVA-mCherry*, *C. glutamicum divIVA::divIVA-mNeonGreen*, and *C. glutamicum rodA::rodA-eYFP* were cultivated in BHI medium (Oxoid) at 30 °C and 200 rpm. Benzothiazinone 043 (Caychem) was dissolved in dimethyl sulfoxide (DMSO) and directly added to a final concentration of 1 µg ml^−1^, if not indicated differently. EMB added in the same way was used at 10 µg ml^−1^. For growth curve analysis, cultures were inoculated with 200 µl of overnight cultures into 10 ml BHI medium and incubated at 30 °C while shaking (200 rpm). Growth curves were plotted based on the optical density (OD_600nm_), measured with an Ultraspec 2000 UV Visible Spectrophotometer (Pharmacia Biotech). Samples for fluorescence and for electron microscopic analyses were taken 3 and 4 hours after inoculation into fresh medium, supplemented with the respective antibiotics, unless otherwise stated. For mycolic acid (MA) analysis, crude cell walls, derived from cell disruption and washing, were further purified with SDS and proteinase K (CW_SDS-PK_), as described before (Braun and Rehn, 1969; Schubert et al., 2017). Cells for cell wall analysis were grown for 3 hours.

**TABLE 1:** *Corynebacterium glutamicum* strains used in this study.

| Name | Genotype | Reference |
| --- | --- | --- |
| *Corynebacterium glutamicum* 13032 | wild type | DSMZ |
| CDC010 | RES 167, *divIVA::divIVA-mCherry* | (Donovan et al., 2012) |
| B4B7 | RES 167, *divIVA::divIVA-mNeonGreen* | (Schubert et al., 2015) |
| BSC026 | RES 167, *rodA::rodA-eYFP* | (Sieger et al., 2013) |

### Fluorescent staining

For all staining protocols, 1 ml of a growing culture was pelleted for 2 min, at 10464 x *g*. For the simultaneous staining of the phospholipid- and the myco-membrane, cells were resuspended in 200 µl phosphate-buffered saline (PBS; 137 mM NaCl, 2.7 mM KCl, 10 mM Na_2_HPO_4_, and 1.8 mM KH_2_PO_4,_ pH 7.4) together with 2 µl of 100 mg ml^−1^ Nile Red (Sigma) and further incubated for 5 min at 30 °C, shaking in the dark. Cells were harvested, washed twice, and finally resuspended in 50 µl PBS. For specific staining of the outer leaflet of the myco-membrane, cells were resuspended in 200 µl PBS together with 2 µl of 10 mg ml^−1^ 6-azido-trehalose (AzTre) (JenaBioscience) and further incubated for 5 min at 30 °C, shaking (Swarts et al., 2012). Cells were harvested, washed twice, and resuspended in 200 µl PBS. After adding 1 µl of 50 mg/ml DBCO-carboxyrhodamine (CR), cells were incubated at RT for 10 min in the dark. Again, cells were washed twice and finally resuspended in 50 µl PBS. For the visualization of nascent peptidoglycan, cells were resuspended in 25 µl PBS together with 0.25 µl of 5 mM 3-[[(7-Hydroxy-2-oxo-2*H*-1-benzopyran-3-yl) carbonyl]amino]-D-alanine hydrocholoride (HADA) (Tocris) dissolved in DMSO, vortexed and briefly spun down (Kuru et al., 2012). After incubation at RT for 5 min in the dark, cells were washed twice and finally resuspended in 50 µl PBS. From each final preparation, 2 µl were used for phase contrast and fluorescence microscopy (Carl Zeiss Axio Imager Microscope) on an agarose pad (1% agarose, Serva) dissolved in ddH_2_O.

### Quantitative MA analysis

For the extraction and esterification of the MA, a modification of a described protocol was used (Goodfellow et al., 1976). 10 mg of purified cell walls (CW_SDS-PK_) were dissolved in a 600 µl of a water-free (30:15:1, vol/vol/vol) mixture of methanol, toluol and 95% sulphuric acid and incubated for 16 h at 50 °C. Further, 400 µl of petroleum ether (boiling point of 60 - 80 °C) were added. After shaking, the petrol ether phase was dried in synthetic air and dissolved in 20 µl of petroleum ether directly before use in the thin layer chromatography (TLC). The TLC was performed on Silica 60 (Merck, Darmstadt) plates with 0.1 mm thickness. As mobile phase a mixture of toluene-acetone (97:3, vol/vol) was used. The development reagent was made of 3 g molybdato-phosphoric acid dissolved in water-isopropanol (25:75, vol/vol). After application of the reagent, the plates were developed at 100 °C for 20 min and further analyzed with the ChemiDoc-System (Bio-Rad, USA).

### Wide-field and fluorescence microscopy

Wide-field microscopy was performed on a Zeiss Axio Imager M1 microscope Plan-Apochromat 100/1.4 oil Ph3 objective (Zeiss). Fluorescence was visualized with the appropriate filter sets (Zeiss). Images were acquired with AxioVisioin (Zeiss) and processed with Fiji (31).

### FRAP

For fluorescence recovery after photobleaching (FRAP) experiments, an LSM 900 Airyscan imaging system (Zeiss) equipped with a Plan-Apochromat 40x/1.4 Oil DIC (UV) VIS-IR M27 objective was used. The bleaching of circular spots was performed with a 488 nm laser at 100% power and a 500 ms pulse. The recovery of carboxyrhodamine labeled azido trehalose was measured every 2.4 s over 40 iterations. The analysis of the half-time recovery was performed as described previously (Giacomelli et al., 2022). The corrected total fluorescence was calculated by measuring the integrated density of a region of interest (ROI), and correcting for background noise and photobleaching.

### Transmission electron microscopy

Ultrastructure of the cell envelope was analyzed on thin resin sections by transmission electron microscopy (TEM). Bacteria were cultured at the exponential phase for 4 hours either in control conditions, or in the presence of 1 µg ml^−1^ BTZ or 10 µg ml^−1^ EMB. For fixation, 2% glutaraldehyde (GA) in 200 mM HEPES buffer, pH 7.4 was added directly to the culture medium at a 1:1 volume ratio. After 10-minutes of initial fixation, bacteria were pelleted, resuspended in 1% GA in 200 mM HEPES buffer and incubated overnight at room temperature with gentle agitation for continued fixation. For resin embedding, cells were post-fixed with 2% osmium tetroxide in 1.5% potassium ferricyanide/dH_2_O for 1 h on ice, followed by 0.2% tannic acid in 100 mM HEPES buffer, pH 7.4 for 15 min. Next, cells were embedded in 1% low-melting point agarose, contrasted en-bloc with 2% aqueous uranyl acetate for 1.5 h at room temperature, and dehydrated with an ethanol series 70-80-90-96-100-100%, each step for minimum 15 min. After progressive infiltration with epoxy resin, samples were polymerized at 70°C for 2 d. Ultra-thin, 80-nm-thin sections were cut using a Leica UC7 ultramicrotome, deposited on copper, slot, formvar-coated grids, and contrasted with saturated aqueous uranyl acetate for 10 min, followed by lead citrate for 3 min. All grids were imaged in a Tecnai G2 Spirit BioTWIN transmission electron microscope (FEI, now Thermo Fisher Scientific), operated at 80 kV, and equipped with a LaB6 filament, an Eagle 4k x 4k CCD camera and TIA software (both FEI, now Thermo Fisher Scientific).

### Scanning electron microscopy

Cells were grown and fixed as described for TEM analysis. A small volume of a dense suspension of glutaraldehyde-fixed bacteria was transferred onto carbon and poly-L-Lysine coated cover slips. After 5 min, cover slips were washed with water, and samples dehydrated with an ethanol series 70-80-90-96-100-100%, each step for minimum 15 min, followed by critical point drying (CPD 030, BAL-TEC AG). Dried cover slips were mounted on adhesive carbon tape and sputter coated with 3 nm Platinum. Samples were imaged in a Sigma 300 VP (Zeiss) scanning electron microscope using a secondary electron (SE) detector and 1 kV accelerating voltage.

### Antibiotic interaction testing

The test for antibiotic interaction was performed as described before, using BHI medium (Lechartier et al., 2012; Palomino et al., 2002; Schubert et al., 2017). In horizontal rows, two-fold serial dilutions of BTZ starting at the MIC were prepared in 96-well microtiter plates. Respectively, two-fold serial dilutions of the antibiotics penicillin G (PenG), carbenicillin (Carb), kanamycin (Kan), ethambutol (EMB), or rifampicin (Rif) were added in the vertical direction, starting at the respective MICs. The first row and column served as negative controls. An exponential-phase *C. glutamicum* ATCC 13032 culture was diluted to OD_600_ of 0.001, and 20 µl were added to each well using a Liquidator96 (Mettler-Toledo) pipette system. The microtiter plate was incubated in a ThermoMixer at 30 °C and 700 rpm for 16 h. 20 µl of resazurin (0.01% in distilled water, filter sterilized) was then added to each well, and incubated in a ThermoMixer at 30 °C and 300 rpm for 30 min. Tests were done in parallel, both in a clear microtiter plate for visual inspection and in a black microtiter plate for fluorescence measurement. Fluorescence of resorufin was recorded in a Tecan Infinite M200 Pro (Tecan) microplate reader (excitation wavelength, 560 nm; emission wavelength, 590 nm). Mean values calculated from two to five experiments are shown in a heat map prepared with R.

### Computational image analysis

For the systematic analysis of fluorescent micrographs, the previously used custom-made solution was further developed (Schubert et al., 2017). Based on the construction of an exact centerline for single cells, a systematic measurement along the outline of cells became possible. By this, a distinct analysis of membrane and cell wall regions can be derived from the same dataset that was used for the analysis of cytosolic signals. The FIJI- and R scripts are available on request.

## Results

### Minimum inhibitory concentration

First, we determined the minimum concentration of BTZ that stopped growth of *C. glutamicum*) based on OD_600_ measurements. For this a wild type strain expressing a DivIVA-mCherry fusion (*C. glutamicum divIVA::divIVA-mCherry*) was used to allow visualization of pole formation and cytokinesis as described before (Schubert et al., 2017). BTZ was added to freshly inoculated cultures. The concentration of 1 µg ml^−1^ (2.3 µM) BTZ was sufficient to stop growth about two hours after the treatment, but did not lead to considerable cell lysis (Fig. 2A). The bacteriostatic inhibition was reversible and cultures recovered after 25 hours (Fig. 2A, Fig. S1, S5). Higher concentrations of BTZ (10 µg ml^−1^ and 50 µg ml^−1^) did not allow recovery of growth within 25 h. As observed with phase contrast microscopy, cells appeared shorter and thicker while an accumulation of the polar scaffold DivIVA-mCherry became visible (Fig. 2B). Accumulation of DivIVA at the cell poles is indicative of a reduced elongation growth as shown before (Schubert et al., 2017).

**Figure 2:**
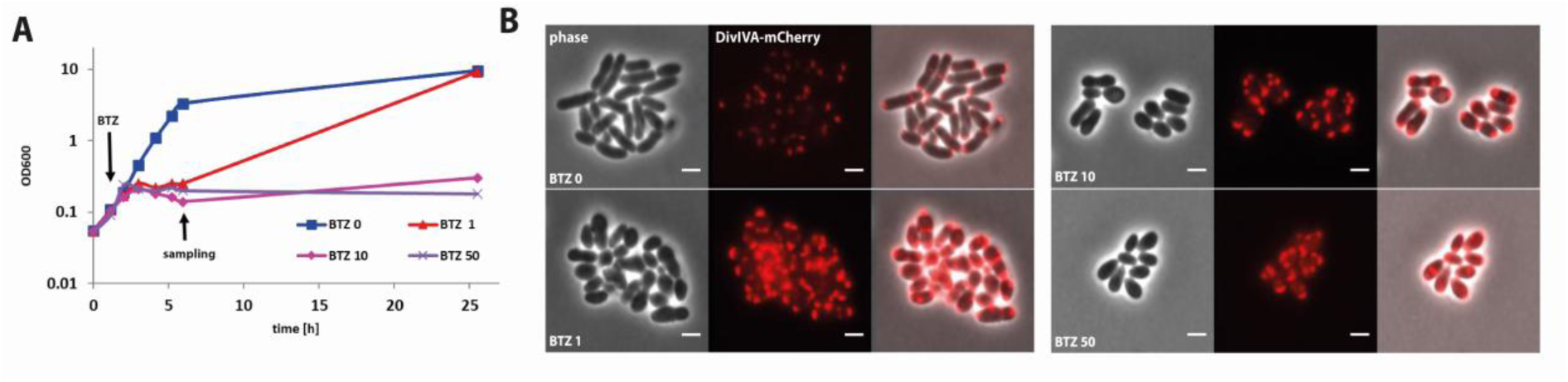
I*n vitro* effects of BTZ043 on *C. glutamicum* growth. **(A)** Addition of BTZ043 to *C. glutamicum* (here a strain carrying an allelic replacement *divIVA::divIVA-mChrerry*) has a bacteriostatic effect. Already 1 µg ml^−1^ BTZ leads to a quick arrest of cell growth, which is reversible, while higher concentrations of BTZ show a prolonged bacteriostatic effect. **(B)** Fluorescence microscopy revealed that BTZ treated cells appear shorter and thicker. The polar scaffold protein DivIVA (shown here as DivIVA-mCherry fusion) accumulate at the cell poles and the division-plane. Scale bar: 2 µm.

Treatments with similar doses of BTZ but applied after three hours of growth (at OD 0.4) also showed a comparable impact on cell shape and influenced growth within two hours. Doses between 1-20 µg ml^−1^ lead to a gradual decrease of the growth rate (Fig. S1A).

### Cell wall synthesis

We have shown previously that EMB treatment leads to a stop of apical PG synthesis and so we next investigated the effects of a sublethal concentration of BTZ on cell elongation and cytokinesis using fluorescence biorthogonal labelling or fluorescent reporter strains (Fig. 3). As part of the image analysis, individual cells were isolated from micrographs and further processed. Here, either the length axis, or the cell outline served as a topological reference (Messelink et al., 2021). This way, in *C. glutamicum,* polar signals originating at old or young cell poles could be distinguished. This allowed for a consequent downward orientation of the old (brighter) pole in the linear single cell data. The resulting set of visualizations showed the cumulative data from microscopic experiments, each sorted by increasing cell lengths, and respectively color coded for the signal intensity.

**Figure 3:**
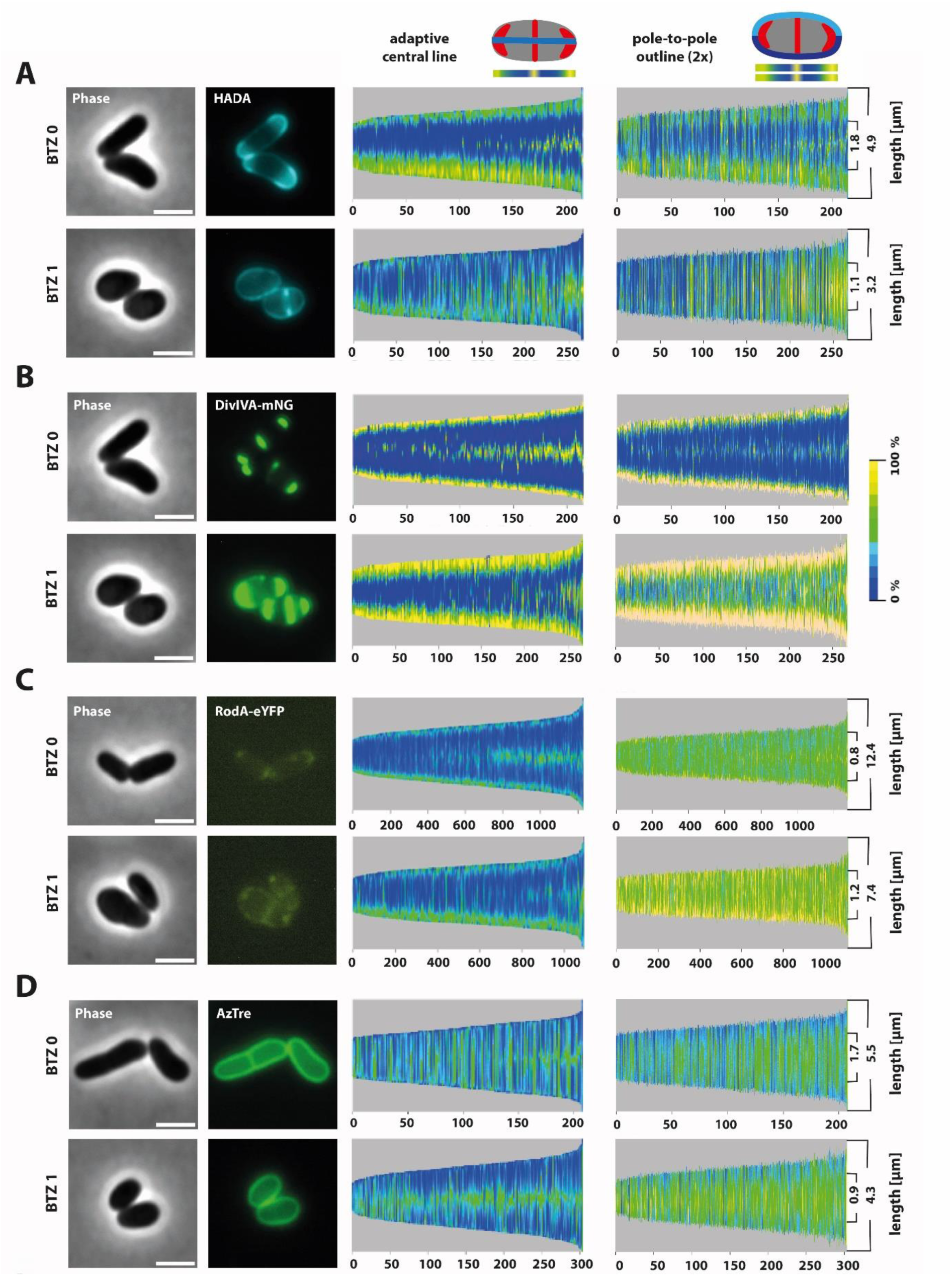
Phenotypic characterization of the BTZ effects on *C. glutamicum* cell wall synthesis. Phase contrast and fluorescence micrographs (left) and demographs derived from multi-cell analysis (right). All experiments were performed with a *C. glutamicum* strain carrying an allelic exchange of the *divIVA* by the *divIV::divIVA-mNeonGreen* fusion allele. **(A)** HADA staining revealed the characteristic pattern of polar and septal PG synthesis including the asymmetric polar synthesis rates. Treatment with BTZ abolishes apical HADA incorporation, indicating a block in the elongation growth. **(B)** The polar scaffold protein DivIVA, visualized here as DivIVA-mNeonGreen fusion, localizes to cell poles and late septa. After BTZ treatment DivIVA accumulates, leading to cell widening and bulging. **(C)** The elongasome specific transglycosidase RodA (shown here as an allelic replacement r*odA::rodA-eYFP*) exclusively localizes to cell poles in untreated cells. After BTZ addition, RodA mislocalizes over the entire membrane. However, most poles retained a focal accumulation of RodA, which becomes explicitly clear in the demographs. **(D)** AzTre labelling of *C. glutamicum* cells indicates the mycolic acids in MM. For both, control and treated cells, a uniform and continuous MM is visible around the whole cell. Scalebar: 2 µm.

We used a fluorescent D-alanine analog (HADA) to stain sites of nascent PG synthesis (Fig. 3A). In untreated cells, HADA signals were localized to the poles, and in dividing cells, to the septum, consistent with earlier reports (Messelink et al., 2021). Addition of BTZ led to a complete loss of the apical cell wall synthesis, similar to the treatment with EMB (Schubert et al., 2017). Not surprisingly, BTZ treated cells were about 1 µm shorter in length, indicating the loss of efficient apical elongation growth (Fig. S1B, Histogram). PG seemed to be synthesized only at sites of cell division and surprisingly some signal was observed along the lateral walls. Therefore, we next analyzed the localization of the polar scaffold protein DivIVA (*C. glutamicum divIVA::divIVA-mNeonGreen*) and the major elongasome specific transglycosylase RodA (*C. glutamicum rodA::rodA-eYFP*). DivIVA is considered a central organization hub for the apical growth and permanently localizes to newly forming poles during cell division (Giacomelli et al., 2022; Letek et al., 2008; Melzer et al., 2018). In untreated cells, DivIVA localized as expected to the old and new cell poles (Fig. 3B). In BTZ treated cells, DivIVA signal was localized predominantly to old cell poles, its drastic accumulation reflects slower elongation growth as shown previously (Schubert et al., 2017). Only a small fraction of cells showed the signal also at new poles, which could suggest that a fraction of cells in the process of septation is reduced. DivIVA interacts with RodA, a key factor for the expansion of the peptidoglycan (PG) sacculus (Emami et al., 2017; Meeske et al., 2016; Sieger et al., 2013; Sieger and Bramkamp, 2015). In untreated cells, RodA was localized exclusively to the cell poles (Fig. 3C). Upon BTZ treatment, RodA localization became more dispersed. It was still localized to old poles, but due to the absence of polar PG-synthesis, indicated by the missing HADA-signal, this pool of RodA was probably inactive. In addition, the RodA signal was localized to young poles and occasionally even to the sidewall. This is consistent with the localization of the HADA signal and indicates that the synthesis of the cell-wall in BTZ treated cells is most intense at new poles but can occur also at the lateral sites.

### Integrity of the MM

We used 6-azido-trehalose (AzTre) bioorthogonal labelling to visualize the outer leaflet of the MM (Fig. 3D) (Swarts et al., 2012). In BTZ treated cells, we did not observe any apparent difference in the fluorescent signal of the AzTre labelling across the cell surface, indicating that BTZ treated cells retained a confluent MM (Fig. 3D). This was surprising compared to the loss of the MM on the cell poles in EMB treated cells (Fig. S2) (Schubert et al., 2017). Obviously, BTZ treated cells are able to cope with the shortage of Ara*f* occurring due to sublethal BTZ treatment in the sense that the MM stays confluent. We proceeded by quantification of total bound MA in crude cell wall preparations using thin layer chromatography (TLC) (Fig. 4A). The crude cell wall samples were prepared by mechanical disruption of cells using a boiling SDS solution and proteinase K treatment. Mycolic acid methyl esters (MAME) were only slightly reduced after BTZ treatment. Again, this effect was relatively mild compared to a significantly decreased MA content in EMB treated cells (Schubert et al., 2017).

**Figure 4:**
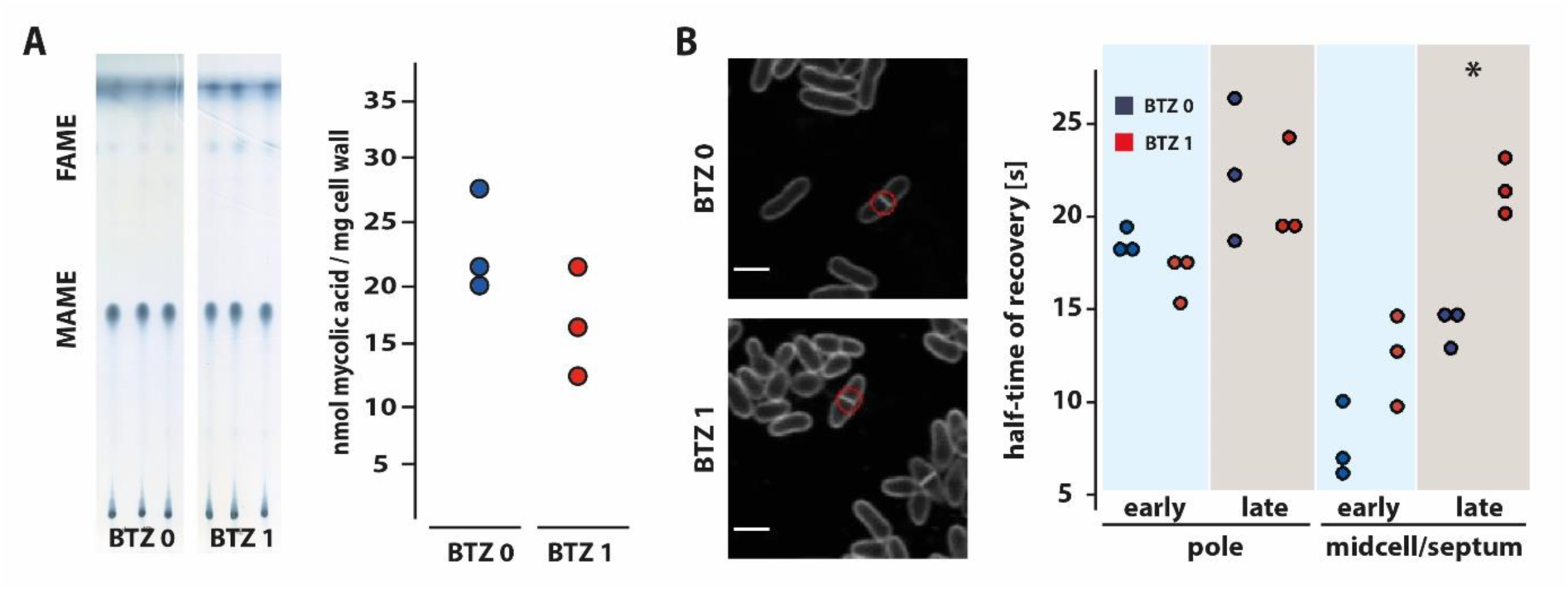
Quantification and dynamics of MM in BTZ treated *C. glutamicum* cells. **(A)** The mycolic acid content, extracted from 1 mg crude cell wall preparation, was analyzed densitrometrically, after running a thin layer chromatography (TLC). No significant difference was observed between the content of mycolic-acid methyl-esters (MAME) (unpaired t-test, p = 0.15). **(B)** Dynamic properties of the MM at polar- and midcell-regions were analyzed with fluorescence recovery after photo-bleaching microscopy (FRAP). A significantly faster recovery (unpaired t-test, p = 0.01) was measured only for the septal signal of treated cells. Scalebar: 2 µm.

With FRAP-microscopic analyses of the AzTre signal, dynamic properties of the MM were examined (Fig. 4B). Recovery of the AzTre signal at polar and mid-cell loci was compared between BTZ treated and untreated control cells, at the early and at the late stage of the cell-cycle, defined by the absence or the presence of a division septum, respectively. At the cell poles, neither the early-stage, nor the late-stage cells showed a significant difference in the half-time of the AzTre signal recovery between treated and control cells. Furthermore, no influence on the dynamics of the MM was observed upon BTZ treatment at the sidewall in the pre-septal stage. In contrast, the fluorescent signal derived from the division-septum showed a significantly slower recovery for BTZ treated cells. This difference could be explained by the impaired *de novo* formation of the underlying scaffold that in BTZ treated cells only occurs at the septum, as shown in Fig. S2.

For further confirmation of the MM integrity, we next analyzed the cell surface by scanning electron microscopy (SEM) (Fig. 5). In control cells, the cell surface appeared smooth and uniform, indicating an intact MM covering the entire cell surface (Fig. 5, upper panel). In BTZ treated cells, the outer layer revealed crevices on the poles and some furrows at the sidewall below. Some excess material was visible on the surface as small vesicles. Despite this apparent destabilization, the MM still covered the entire cell surface, including the cell poles. However, occasionally a pair of hinged daughter cells was observed whose emerging young cell poles did not seem to be covered with the MM (Fig. 5, middle panel). Consistent with our earlier report (Schubert et al., 2017), and contrary to the effects of BTZ, in cells treated with EMB the loss of confluency in the MM was striking. We observed wide furrows that extended longitudinally across the cell surface, and crevices over the cell poles. Young poles of hinged cells seemed to lack the outermost layer entirely (Fig. 5., lower panel).

**Figure 5:**
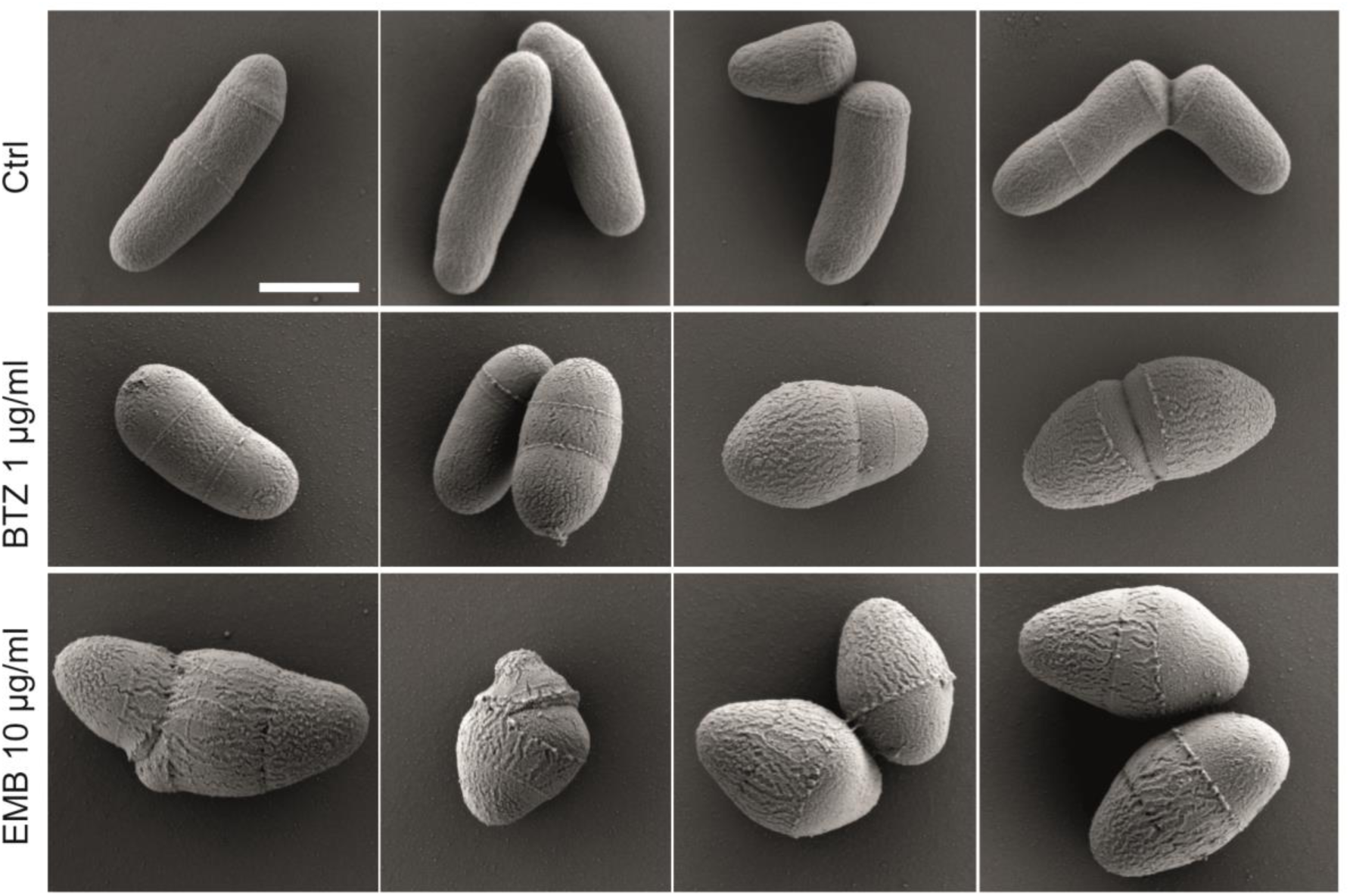
Scanning electron microscopy of untreated and of BTZ or EMB treated *C. glutamicum* cells. Control cells (upper panel) reveal a smooth cell surface and rod-shaped morphology. Cells treated with BTZ (middle panel) are shorter and thicker compared to untreated cells and occasionally have excess material blebbing from the surface. Furrows originating from the cell poles are visible, indicating fractures in the outer envelope layers. Emerging new poles sometimes seem to lack the outer layer. Cells treated with EMB (bottom panel) have ovo-coccoidal shape and wide furrows all over the cell surface. The new poles after cell separation seem to lack completely the outer layer, similar to data published before (Schubert et al., 2017). Scale bar: 1 μm.

### Cell morphology

In addition to the confluency of the MM, SEM images revealed considerable morphological differences between control and treated cells that are likely related to the changes in the expansion of the cell wall. Whereas control cells were rod-shaped, BTZ treated cells became shorter and thicker. In comparison to EMB treated cells, most of BTZ treated cells were not obviously pointed and the frequency of hinged daughter cells was lower (Fig. 5, Fig. S3). Moreover, in control cells, it was very difficult to find daughter cells in the process of daughter cell separation. We have observed only a couple of cells with a shallow groove in the cell surface perpendicular to the longitudinal axis, or with tiny perforations on the cell surface above the septum (Fig. 5, Fig. S4), as reported previously (Zhou et al., 2019). In contrast, separation intermediates were more frequent in BTZ and in particular in EMB treated cells (Fig. S3). They were represented by single cells with perforations on the cell surface, or by attached daughter cells only partially separated as evidenced by partially exposed young cell poles (Fig. S4). This indicates that the process of daughter cell separation in treated cells was slower. A growth curve performed for both antibiotics in parallel revealed that BTZ treated cells grew slower than EMB treated cells (Fig. S5).

### Ultrastructure of the cell envelope

We next performed ultrastructural TEM analysis to visualize the cell envelope in cross-sections (Fig. 6). Samples were prepared conventionally by chemical fixation and epon resin-embedding. In the untreated control cells, multiple layers of distinct electron densities could be resolved. We assigned them molecular identities based on previous studies (Zhou et al., 2019; Zuber et al., 2008). At the lateral side (Fig. 6, top left panel), the innermost cytoplasmic membrane was visualized as two layers; an electron translucent layer was assigned to phospholipid tails, and a thin electron dense layer above it was assigned to phospholipid heads in the outer leaflet. The layer of phospholipid heads in the inner leaflet was not well resolved because of the intense staining of the cytoplasm. The less electron dense layer above the cytoplasmic membrane is considered to be the periplasmic space (PP). This was followed by a thick electron dense layer, which consisted of the PG with the associated AG. The surface electron translucent layer corresponded to the MM. The outermost electron dense line likely depicts the hydrophilic trehalose head-groups of the outer MM leaflet, made of TMM and TDM. In BTZ treated cells, the overall thickness of the cell envelope was similar to control cells, but there were some structural changes (Fig. 6, middle panel). The periplasmic space was not discernible, while the electron dense layer of the outer leaflet of the cytoplasmic membrane appeared thicker. We hypothesize that a DPR precursor that is predicted to accumulate in the periplasm upon BTZ treatment (Grover et al., 2014) increases its electron density so that the outer leaflet of the cytoplasmic membrane and the periplasm appear as one layer. Interestingly, this electron dense layer was notable also at the lateral sides, in places where active cell wall synthesis does not normally take place. The PG/AG layer did not appear notably different. It was covered with the MM, which formed a continuous layer although it occasionally appeared less regular than in untreated cells (Fig. S6), likely reflecting the furrows that were observed by SEM. Differences between control and BTZ treated cells were more prominent along the forming division septum (Fig. 6., middle panel). Whereas in control cells, a division plane was not visible on the cell surface, in BTZ treated cells the position of a septum was revealed by a shallow groove. The increased radius at the junction between the lateral side and the division septum in BTZ treated cells (Fig. 6., white arrows) demonstrates that the lateral PG layer has less mechanical resistance to the cytoplasmic turgor compared to untreated cells. In control cells, a Π-shaped structure was formed at the T-junction by two thin, electron dense layers of PG/AG extending into the septum from the lateral cell envelope (Fig. 6, left panel). These two septal PG/AG layers were separated by a layer of low electron density whose composition is unknown. We hypothesize that it is made of MA covalently linked to the AG and thus forming the inner leaflet of the mature MM, which we also refer to as the greasy slide. But until after the snapping division they do not seem to be covered with the outer layer of the MM, including the TMM and TDM. In BTZ treated cells, neither the Π-structure nor the middle layer made of MA were apparent.

**Figure 6:**
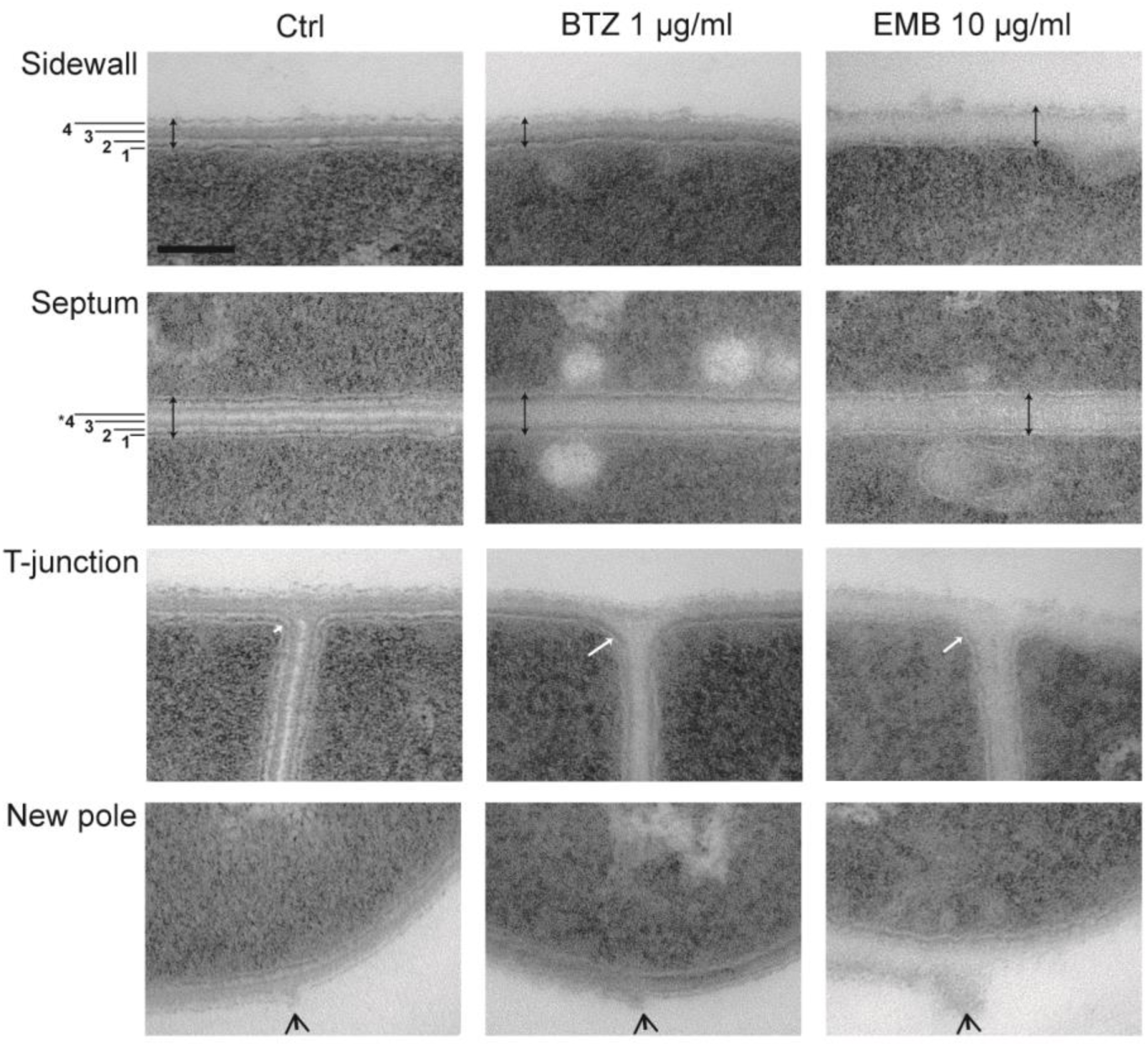
Ultrastructural analysis of untreated and of BTZ or EMB treated *C. glutamicum* cells. In control cells, four distinct layers can be resolved at the sidewall: (1) the cytoplasmic membrane (PM) (2) the electron translucent periplasmic space (PP), (3) a thick and electron dense layer of the peptidoglycan (PG) and the attached arabinogalactan (AG), and (4) the electron translucent MM (MM). In BTZ treated cells, the electron translucent PP is missing, but an electron dense layer is visible in its place. In EMB treated cells, the envelope is thicker. The PG layer is electron translucent and topped with a thin electron dense layer, possibly corresponding to the AG layer in control cells. At the septum, two sets of the first three layers are mirrored across the joint MM layer, which we propose corresponds to the inner MM leaflet only (*4). This layer is not distinct in treated cells. Double sided vertical arrows indicate thickness of the envelope. White arrows indicate the radius between the lateral and the septal cell-wall. Arrowheads indicate birth scars, the remains of what used to be junctional PG that stretched across the septum. Scale bar: 100 nm.

Surprisingly, the cell envelope in EMB treated cells differed considerably from BTZ treated cells in that it was thicker, but its middle layer, likely corresponding to the PG, had reduced electron density. The electron dense layer that was observed in BTZ treated cells below the PG and is considered to be the periplasm filled with DP-linked precursors was not present. Instead, electron dense material was observed between the PG and the MM, and because of its position, we propose that it represents the galactan backbone now displaying a stronger contrast than the PG (Fig. 6, right panel). The septum often bulged into the cytoplasm of one daughter cell (Fig. S6). This indicates that newly built septa have low mechanical strength and that cell wall synthesis at the septum continues well after septation is completed, while daughter cells separation is delayed. Similar to BTZ treated cells, the Π-shaped structure was not formed at the T-junction. The putative low contrast PG layer in the lateral side extended into the septum, but the translucent middle layer, likely made of MA, was not apparent (Fig. 6., middle panels). As a unique feature of EMB treated cells, a thin, electron denser line in the middle of the septum could sometimes be observed. It seemed to be of the same origin as the peripheral electron dense layer observed between the PG and the MM, thus corresponding to a degenerated AG layer.

We carefully compared the appearance of the MM in the three samples (Fig. S6). In control cells, the MM was smooth and continuous all around the cell. In BTZ treated cells, the MM was also continuous, except at the cell poles where the surface appeared rough and the MM could often not be resolved. Young poles of hinged cells were mostly covered with the MM. This was again in stark contrast to the EMB treated cells, where the MM was discontinuous all over the cell surface and completely missing over the young poles of attached cells. Furthermore, it often appeared that furrows in the cell envelope extended below the MM.

### Antibiotic synergy testing

Because results overwhelmingly indicated that the outer leaflet of the MM remained largely confluent after the BTZ treatment, we next tested for the synergistic effects of different antibiotics (Fig. 7.) using the REMA checkerboard assay (Palomino et al., 2002). The in vitro combination of BTZ with several other antibiotics has been tested before in *M. tuberculosis* (Lechartier et al., 2012). We were interested to observe if the effect in *C. glutamicum* is similar. For the β-lactam antibiotics carbenicillin and penicillin G, the test only showed additive effects. Similarly, only additive effect was obtained for the combination with EMB. Rifampicin, a drug commonly used for TB treatment served as a control for a hydrophobic compound and resulted in an indifferent effect, similar to effects observed with kanamycin. These results show that although BTZ had a strong effect on cell morphology and even on the cell wall ultrastructure of *C. glutamicum*, it did not reveal any synergistic effects with the tested antibiotics, in contrast to the EMB treatment (Schubert et al., 2017).

**Figure 7:**
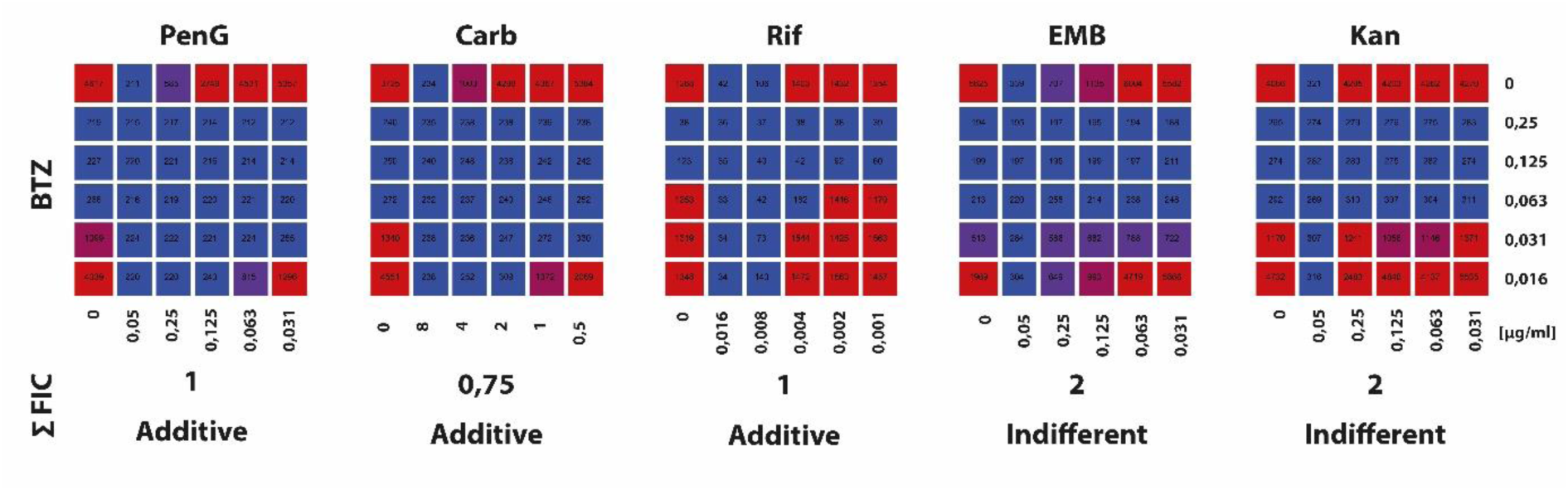
Checkerboard assay for BTZ antibiotic synergy testing in *C. glutamicum*. Using a checkerboard assay (Palomino et al., 2002), BTZ showed additive effects for the β-lactam antibiotics carbenicillin, penicillin G, and for the translation inhibitor rifampicin. For ethambutol and kanamycin, the effect was classified as indifferent.

## Discussion

The synthesis of the complex cell wall of mycobacteria and corynebacteria is an ideal target for antibiotics (Abrahams and Besra, 2018). EMB and BTZ antibiotics were initially both described to interfere with the synthesis of the AG layer and were consequently expected to affect the integrity of a protective MM layer. Our work shows that the effects of the two antibiotics are more complex and partially distinct, which is relevant for consideration about their drug potential. The observations also stimulate reflection on the structure of the cell envelope and the mechanisms of cell division.

The most striking difference between the two antibiotics was that the integrity of the MM was severely impaired after EMB treatment, but considerably less after the BTZ treatment. In EMB cells, the MM layer was absent from young poles, whereas longitudinal furrows interrupted the confluency of the MM across the rest of the cell surface, which is consistent with our previous study (Schubert et al., 2017). In BTZ treated cells, most poles were covered with the MM, but we did observe emerging young poles without the MM. In contrast, the cell surface in control cells seemed uniformly covered with the MM. Recently, assembly of the MM during cells division was elucidated by metabolic staining and it was proposed that the MM of *C. glutamicum* becomes confluent just before V-snapping by the inflow of peripheral trehalose glycolipids into the septum (Zhou et al., 2019). Our observations with fluorescently labelled trehalose similarly suggest that the MM is assembled before V-snapping occurs. Metabolic labelling with AzTre even enabled us to compare dynamic properties of the MM, and the recovery of the septal MM was slower in BTZ treated cells. However, our SEM and TEM analyses did not corroborate the observation that the outer leaflet of the MM would flow into the septum before daughter cells are separated. In control cells, we observed an electron translucent layer in the middle of the septum, between the two septal PG layers. We propose that this layer corresponds to the inner leaflet of the MM, made of MA covalently anchored to the underlying AG.

While the septal PG was continuous with the peripheral PG, this electron translucent layer was isolated from the peripheral MM by the junctional PG. The layer can be observed as soon as the septum starts growing, indicating that all layers of the cell envelope are built synchronously (Fig. S6). We further cautiously propose that in control cells the spreading of the MM outer leaflet across the immobilized greasy slide of freshly exposed young poles would be practically instantaneous, and that the inflow of glycolipids into the septum before daughter cell separation would be of no advantage. We did not observe the electron translucent layer in the middle of the septum in either antibiotic, which argues that the inner leaflet of the MM is a lot thinner if at all present. As a consequence, the lateral flow of TMM, TDM or free MA across the bacterial surface would be severely impaired, and giving rise to the phenotype we have described. Because the MM in BTZ treated cells seemed to be affected less, we speculate that at the sublethal concentration of BTZ, cells maintain ability to build a confluent though less dense layer of covalently attached MA, while the slow process of daughter cell separation may give the MA in the outer leaflet relatively more time to spread over slowly emerging young poles. BTZ blocks the epimerization of DPR to DPA, a lipid-linked agarose precursor that is used for the synthesis of the agarose polymer (Crellin et al., 2011). Deletion of *aftB*, an enzyme involved in the terminal Ara*f* coupling on the arabinan domain, produced a similar phenotype to BTZ. The MM was formed but it was prone to mechanical stress, which can be explained by a reduced number of mycolated sites on the AG (Raad et al., 2010). EMB acts downstream and appears to block enzymes that polymerize branched arabinan. *M. tuberculosis* strains resistant to EMB carry mutations in the arabinosyl transferases EmbABC (Forbes et al., 1962; Sreevatsan et al., 1997; Telenti et al., 1993). Moreover, overexpression of the single EmbC gene in *C. glutamicum* decreased the sensitivity of these cells to EMB (Radmacher et al., 2005). Of note, we observed an electron dense layer immediately below the MM layer on the lateral surface and occasionally in the middle of the septum only in EMB cells. It likely corresponds to the galactan backbone and possibly some additional arabinose sugars. Its irregular appearance suggests that it is structurally altered and may not provide a correct scaffold for covalent attachment of MA. Furrows visible across the surface of the EMB cells in particular are also a hallmark of stalled lateral flow of the MM.

In MTB cells, chemical modifications of MA are differently regulated in active growth and growth stasis (Dulberger et al., 2020). Our chemical analysis revealed that the overall content of MA is slightly reduced after BTZ treatment. We cannot exclude that slowly growing BTZ treated cells produce more MA than faster growing EMB treated cells. Also, in the *aftB* mutant membrane fragments were observed shedding to the external culture medium (Raad et al., 2010).

Changes in the shape of treated cells demonstrate that the effects of the two antibiotics are not limited to the confluency of the MM. Pointed shape can be explained by inhibition of the apical elongation, but continued growth at the division septum, which was observed in both antibiotics by metabolic labelling and localization of fluorescently labelled DivIVA and RodA. But there were also obvious differences between the shape of BTZ and EMB treated cells. The EMB treated cells were notably more pointed. This could be explained by the observation that PG synthesis in BTZ treated was localized not only to the septum/young poles but also to the sidewall, which was not the case in EMB treated cells (Schubert et al., 2017). Therefore, it seems probable that BTZ treated cells can elongate along the sidewall. In contrast, EMB treated cells grow only at the septum. This explains why cells became so wide and why the surface of young poles was so large. In addition, quantitative analysis of fluorescence micrographs presented by demographs showed that a fraction of cells with the DivIVA protein or with new PG synthesis localized at the septum was in EMB treated cells a lot higher, and in BTZ treated cells a lot smaller than in control cells. Together with the growth curve these results demonstrate that growth was stopped more strongly by EMB (10 µg ml^−1^ or 48.9 µM) than by BTZ (1 µg ml^−1^ or 2.3 µM), although its sublethal molar concentration was 20x lower. The exceptional potency of BTZ was observed before and attributed to covalent inactivation of the DprE1 enzyme by BTZ (Trefzer et al., 2012).

Moreover, TEM ultrastructural analysis revealed major differences between BTZ and EMB treated cells in the structure of the cell envelope. The PG layer in EMB cells appeared thicker and less electron dense, which is consistent with a recent report that EMB inhibits also the glutamate racemase (MurI) (Pawar et al., 2019). This would imply that the PG layer is poorly cross-linked and has lower mechanical resistance. In contrast, in BTZ treated cells the PG layer does not appear structurally significantly altered. However, an electron dense layer was apparent below the PG where in control cells the electron translucent periplasm was observed. Corynebacteria use the DP lipid to flip the arabinose precursor from the cytosol to the extracellular site. It was suggested that inhibiting the epimerase reaction by BTZ would block the efficient recycling of the DP molecule (Grover et al., 2014) and that this would lead to a shortage of DP needed for the generation of the PG precursor lipid-II. We propose that increased electron dense layer below the PG in BTZ treated cells represents the periplasm, in which lipid-linked precursors have accumulated because of the DprE1 inhibition.

As demonstrated by TEM images, in control cells the position of a growing septum is not visible on the bacterial surface because the lateral cell envelope stretching over the septum remains straight. In contrast, in BTZ and EMB treated cells the position of the growing septum is revealed soon after septation starts because the lateral sides along the septum bulge out or are torn open. This likely suggests that the mechanical resistance of the peripheral cell envelope is cells treated with antibiotics is reduced. This applies also to BTZ treated cells, although as discussed above the structure of PG did not appear obviously structurally altered. Perhaps AG and LAM, whose synthesis is also affected with these antibiotics, contribute to reduced mechanical resistance of the cell envelope. It might also be plausible that the Π-structure is weakened and thus not prevents deformation at the septal side. Moreover, SEM images show longitudinal furrows and crevices across the surface of treated cells that give the appearance of stretch marks. This characteristic may be a consequence of reduced fluidity of the MM layer and exaggerated by reduced mechanical resistance of the cell wall to radial expansion. In support of this, TEM images of EMB cells suggest that surface furrows often extend below the MM layer, although we cannot exclude that some preservation artifacts were introduced by sample preparation.

With SEM, control bacteria appeared either as single rods, or as hinged cells that remained connected at one side. The latter correspond to daughter cells with freshly exposed young poles after V-snapping. Very rarely did we observe intermediate stages of daughter cell separation. This indicates that the process of cell separation in control cells is very fast, consistent with previous reports that V-snapping takes a few milliseconds only (Zhou et al., 2019, 2016). In contrast, intermediate stages of daughter cell separation were common in BTZ treated cells and in particular in EMB cells, demonstrating that cell separation is slower. Moreover, cells with several septa or septa bulging into one of the daughter cells were commonly observed after EMB treatments. These observations stimulate speculation on the mechanism of cell separation. TEM images show that the two septal PG layers forming the Π-structure are thinner than the peripheral PG. It is therefore reasonable to propose that the turgor force inside daughter cells is able to bulge the thinner septal PG outwards as soon as junctional PG becomes partially degraded and that the newly emerged poles thereby drive daughter cell separation. This is consistent with the mechanism of the mechanical fracture proposed previously (Zhou et al., 2019). In BTZ and EMB treated cells, where the lateral wall along the septum bulges out, the turgor force directed against the new septum is reduced. In EMB cells where the septum of one daughter cell bulged into the other, the turgor force may even be diminished to such extent that the separation is stalled. Such a situation would resemble laser ablation of one of the daughter cells (Zhou et al., 2019). Continued growth at the young poles could also provide force for cell separation. Because of decreased growth in EMB and BTZ treated cells, such force would be produced over a longer period and thus delay cell separation.

Understanding effects of antibiotics on bacterial cell biology is important in medical settings with combinatorial antibiotic treatment regimes. The nontuberculous mycobacterium *M. abcessus* is an emerging pathogen associated with hard-to-treat lung infections. *M. abcessus* is naturally resistant to EMB due to a gene polymorphism in EmbB (Alcaide et al., 1997). It was recently shown that *M. abcessus* is sensitive to DprE1 inhibitor OPC-167832 (Sarathy et al., 2022). A combination of OPC-167832 and the β-lactam antibiotic cefoxitin was also shown to be additive in drug interaction assays for *M. abcessus*, indicating that blocking of DprE1 has likely similar cellular consequences in all mycolata.

In the presented work we have used sublethal concentrations of BTZ and EMB antibiotics to decipher their complex effects on cell biology of *C. glutamicum*. The actions of the two antibiotics differ mechanistically in the sense that BTZ primarily produces a shortage of building blocks for the biosynthesis of the cell envelope, whereas EMB targets cross-linking of building blocks. We have shown that BTZ strongly inhibits cell growth, but affects the confluency of the MM layer to a smaller extent and only has an additive effect in antibiotic synergy testing. Furthermore, septum formation was markedly reduced. This is strikingly different from the effects exerted by EMB. We propose that destructive potential of BTZ for the structure of the cell envelope is diminished by its strong inhibition of cell growth. In contrast, EMB treated cells continue to grow slowly, but build a structurally defective cell envelope and thereby become increasingly sensitive to β-lactam antibiotics.

## CRediT authorship contribution statement

**Fabian M. Meyer**: Conceptualization, Methodology, Investigation, Formal analysis, Writing – original draft. **Urška Repnik**: Conceptualization, Methodology, Investigation, Formal analysis, Writing – original draft. **Ekaterina Karnaukhova**: Methodology, Investigation, Formal analysis. **Karin Schubert**: Conceptualization, Methodology, Investigation, Supervision, Formal analysis. **Marc Bramkamp**: Conceptualization, Supervision, Funding acquisition, Writing – original draft.

## Acknowledgements

We are thanking Laura Lindenthal, Manja Lukner, and Gerhard Wanner (LMU Munich) for help during the initial phases of this work. We kindly acknowledge the Central Microscopy facility staff at Kiel University for expert technical support.

