## Supplemental Material for "Effects of benzothiazinone and ethambutol on the integrity of the corynebacterial cell envelope"

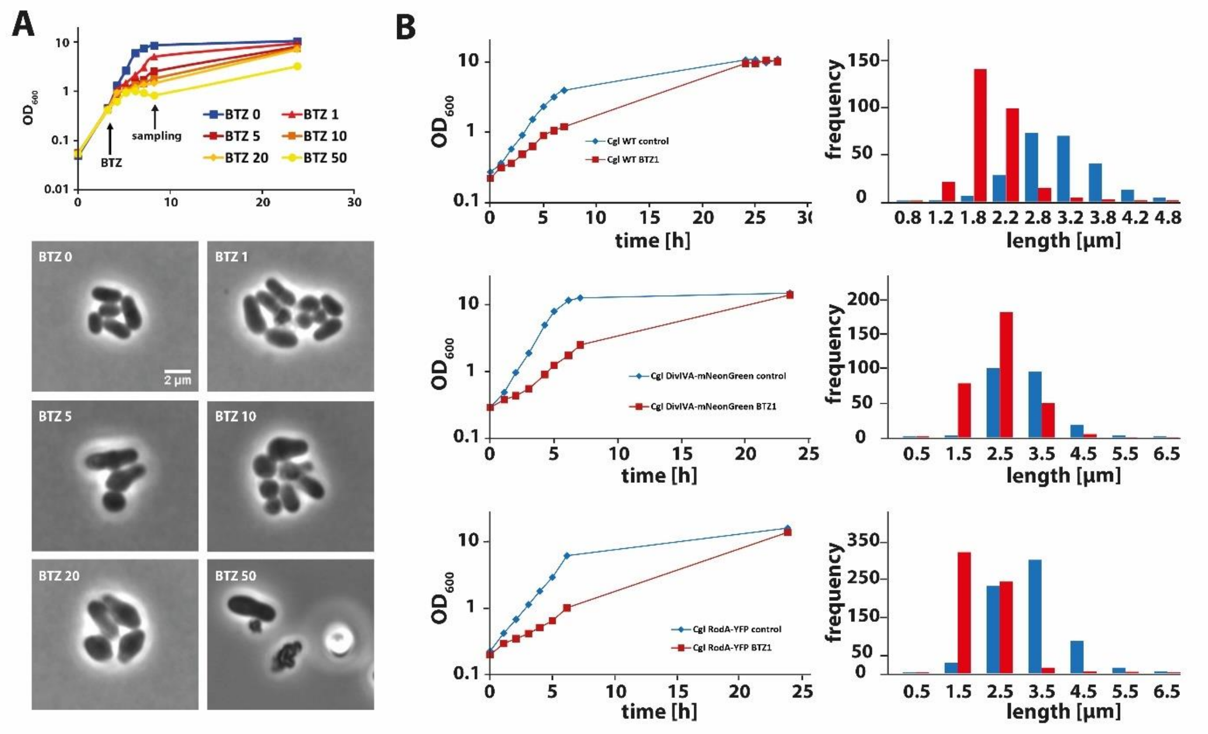


**Figure S1: Additional growth curves and histograms. (A)** Cell growth when BTZ (1 µg $ml^{-1}$ to 50 µg $ml^{-1}$) was added to cultures only after they completed tree cell cycles (OD 0.4). The inoculum expanded roughly by the factor of 10. Recovery within 24 hours was detected for all the applied concentrations, except for 50 µg $ml^{-1}$, meaning that the pool of BTZ was still not fully metabolized. Under the microscope, changes in cell morphology were comparable to experiments with freshly inoculated cells. **(B)** Growth curves and cell length distribution plots in cultures that were used for the fluorescence microscopic analysis, presented in Fig 3. Upon inoculation into fresh medium with 1 µg $ml^{-1}$ BTZ, treated cells of all three strains showed a similar decrease in growth rate. Scalebar: 2 µm.

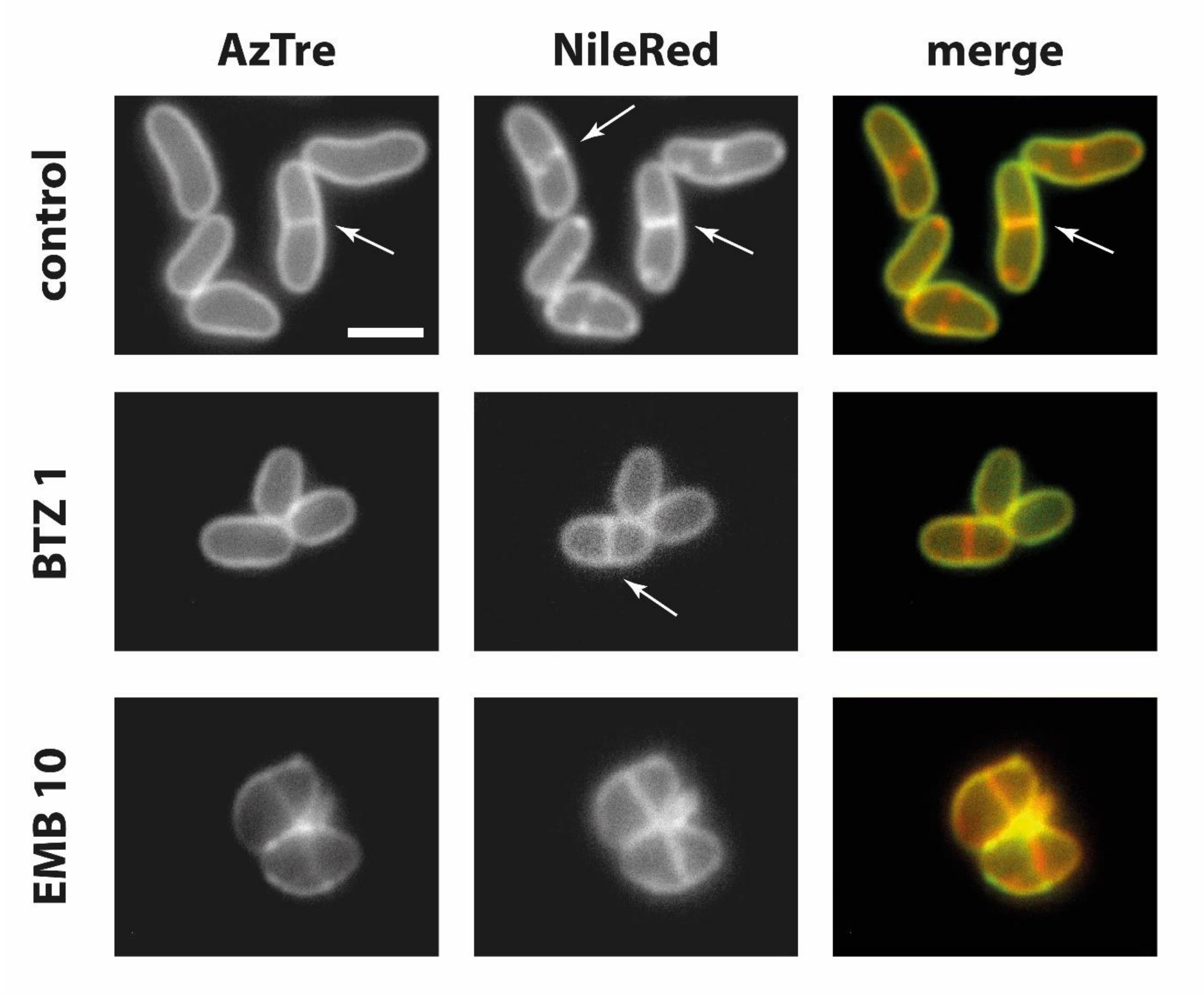


**Figure S2: Effects BTZ and EMB on the MM.** The comparison shows control cells and cells treated with BTZ [1 µg $ml^{-1}$] or EMB [10 µg$ml^{-1}$]. Cells were stained with AzTre to label the MM and with Nile Red to label the PM. Arrows indicate different stages of the septum formation. Both antibiotics result in shorter cells, while central bulging and lemon-shaped poles are most prominent upon EMB treatment. EMB treatment also leads to decreased density of mycolic acids at cell poles. In control cells, as well as in BTZ treated cells, a continuous layer of AzTre labelled mycolates is visible. Note that EMB treated cells show labelling of septa with AzTre before cell separation. Scalebar: 2 µm.

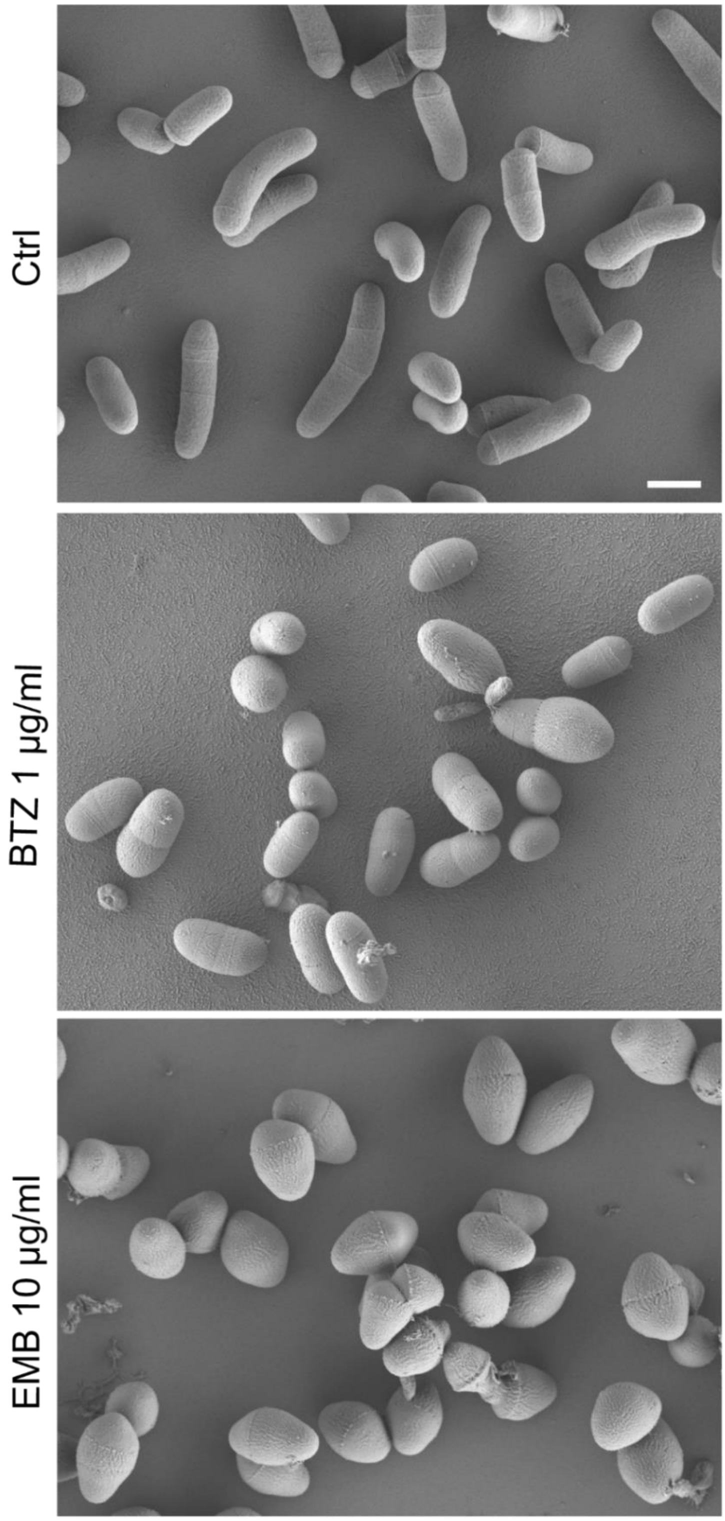


**Figure S3: Cell shape of untreated and of BTZ or EMB treated *C. glutamicum* cells by scanning electron microscopy.** Control cells are rod-shaped, and hinged cells (i.e. daughter cells after V-snapping) are common. BTZ cells are shorter and thicker, while there is notably less hinged cells. EMB treated cells are short and pointed (ovococcoid); hinged cells are frequent. Scale bar: 1 µm.

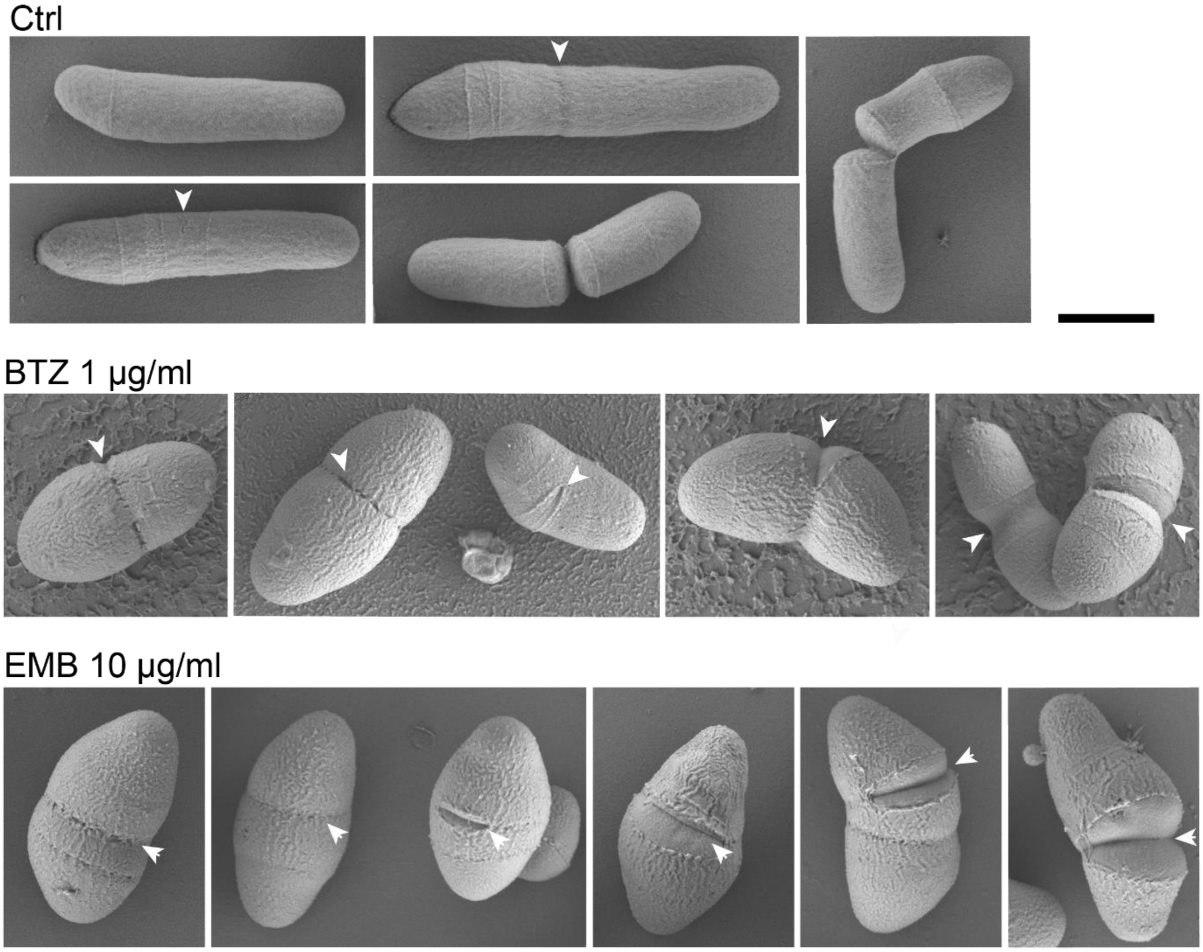


**Figure S4: Intermediate stages of daughter cell separation in untreated and in BTZ or EMB treated *C. glutamicum* cells by scanning electron microscopy.** Control cells (upper panels) appear either as single rod-shaped cells or as hinged cells with exposed young poles. Intermediate stages of daughter cell separation are rare, and are represented by a shallow groove in the cell surface perpendicular to the longitudinal axis, or by tiny perforations on the cell surface above the septum (arrows). Daughter cell separation intermediates are more frequent in BTZ (mid panels) and in particular in EMB (bottom panels) treated cells. They are represented as perforations on the cell surface, or by partially exposed young cell poles (arrows). Scale bar: 1 µm.

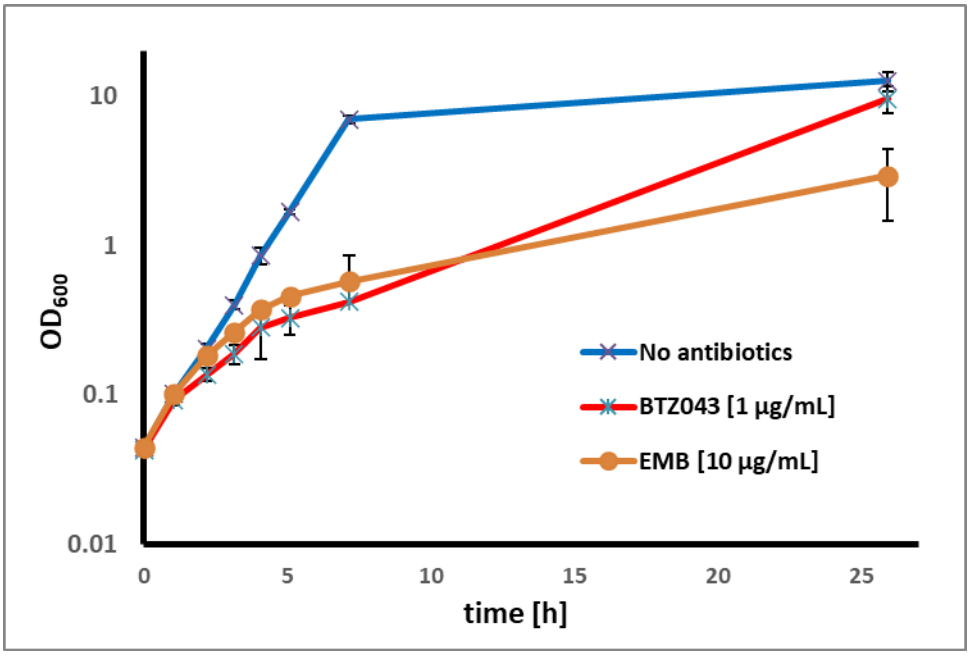


**Figure S5: Effect of BTZ and EMB on the growth of *C. glutamicum* RES167 cells.** The curves represent the mean of biological triplicates including the standard deviation.

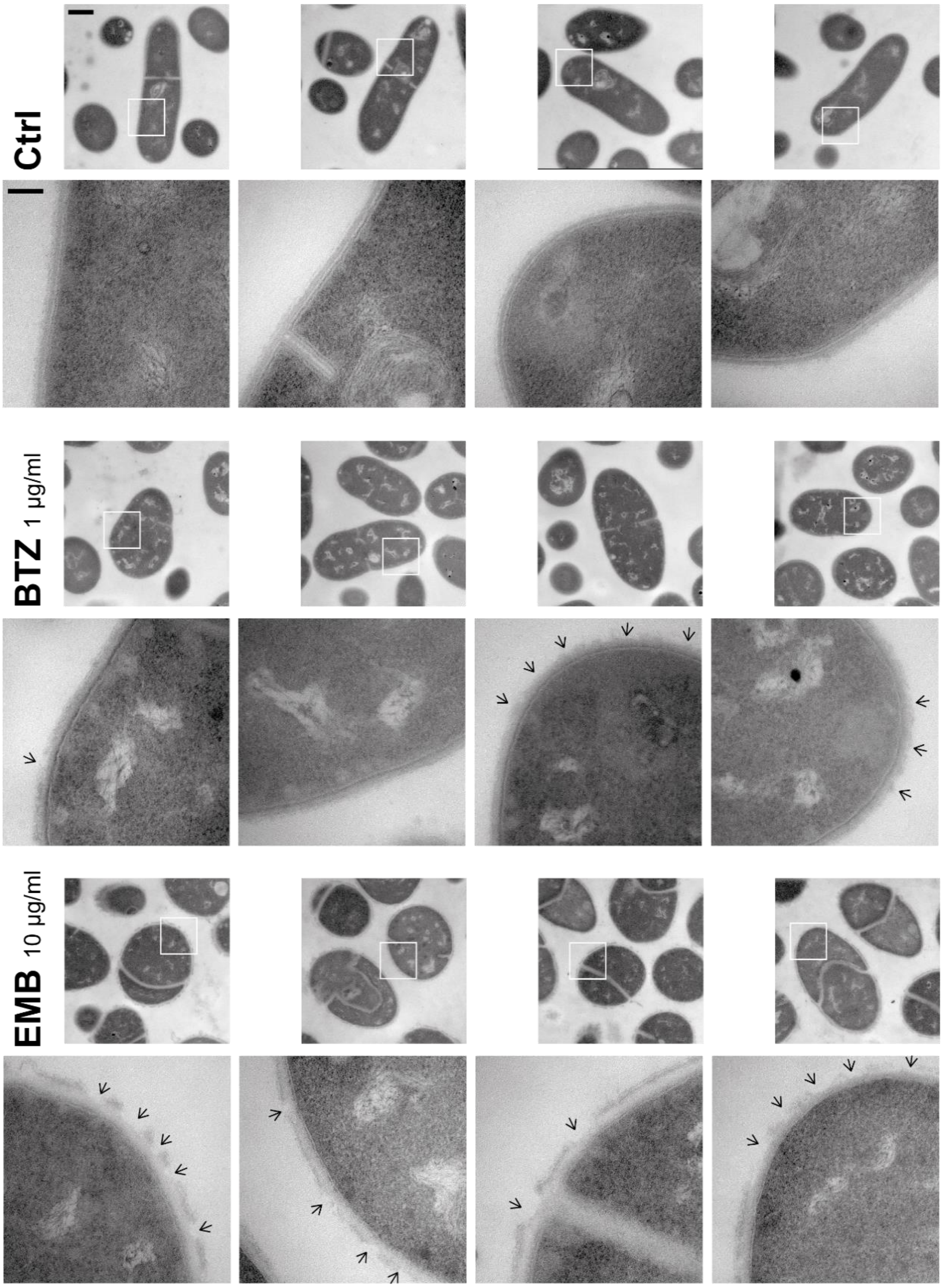


**Fig. S6. Integrity of the MM in untreated and in BTZ or EMB treated *C. glutamicum* cells by transmission electron microscopy.** In control cells, the cell surface is uniformly covered with electron translucent MM. The layers (1), (2), (3) and (*4) (as defined in Fig. 6) can be observed in the growing septum, indicating that layers are built synchronously. In BTZ treated bacteria, the MM does not make an entirely smooth layer, in particular on cell poles, whereas in EMB treated bacteria perturbations in the MM appear all over the surface and often coincide with erosions in the thick electron translucent layer, considered to be the PG. In EMB cells, septa often bulge into one of the daughter cells. White squares indicate regions shown below at increased resolution. Arrows indicate discontinuities in the MM. Scalebar: low mag: 500 nm, high mag: 100 nm.
